# Cholinergic forebrain activation improves cognition, boosts neurotrophin receptors, and lowers Aβ_42_ levels in the cerebral cortex of 5xFAD mice

**DOI:** 10.1101/2022.03.04.482983

**Authors:** Jacob Kumro, Ashutosh Tripathi, Yun Lei, Jeremy Sword, Patrick Callahan, Alvin Terry, Xin-yun Lu, Sergei Kirov, Anilkumar Pillai, David T Blake

## Abstract

The biological basis of Alzheimer’s dementia has been hypothesized in terms of basal forebrain cholinergic decline, and in terms reflecting the neuropathologies surrounding beta amyloid and tau. To shed light on the contributions of these different biological elements, we performed five month intermittent stimulation of the basal forebrain, which projects to the cortical mantle, in 5xFAD Alzheimer’s model mice and wild-type littermates, and subjected mice to behavioral testing and tissue analysis. The 5xFAD mice performed worse in water maze testing than their littermates. Stimulation improved both classes of mice, and removed performance differences between genotypes by the fifth testing day. Stimulated groups had two to four-fold increases in immunoblot measures of each of the neurotrophin receptors tropomyosin receptor kinase A and B. Stimulation also led to lower tissue Aβ_42_ and beta amyloid cleavage enzyme 1 in 5xFAD mice. Despite the lack of strong direct projections from the basal forebrain stimulation region to the hippocampus, the hippocampal tissues in stimulated mice had more nerve growth factor receptor, and lower levels of beta secretase for amyloid. These data support a causal relation between forebrain cholinergic pathways and cognitive decline dependent on Aβ_42_. Activation of cholinergic brain pathways improves neurotrophin pathways and reduces Aβ_42_ accumulation. The recruitment of both classes of neurotrophin receptors in the process suggests a serine protease cleavage intermediary between cholinergic response and neurotrophin activation.

## Introduction

Alzheimer’s dementia was first identified by Alois Alzheimer in the early 1900s as a clear cognitive dementia which resulted in death 4.5 years later. Examination of the brain revealed a diffuse atrophy, particularly in the cortex(Möller & Graeber, 1998). This dementia was later called Alzheimer’s. By the middle of the 1970’s, senile dementia began to be considered of the same origins as familial Alzheimer’s dementia(Terry & Davies, 1980), and studies began to show that deficits in the brain’s cholinergic systems were prominent in the disorder(Coyle et al., 1983; Perry et al., 1978). These hypotheses have stood as the cholinergic hypothesis(Sarter & Bruno, 1997; Terry Jr. & Buccafusco, 2003), and the primary treatment for Alzheimer’s dementia, early in the disorder, are cholinesterase inhibitors(Birks, 2006) that boost the levels of acetylcholine by suppressing its breakdown. The other prominent hypotheses on Alzheimer’s dementia focus on the neuropathological products, beta amyloid plaques and tau neurofibrillary tangles(H. Braak & Braak, 1997; Heiko Braak et al., 2006). While beta amyloid neuropathology is necessary for mild cognitive impairment to convert to Alzheimer’s dementia(Iaccarino et al., 2017), their precise role remains a subject of debate. More recent evidence favors soluble beta amyloid oligomers as being involved in the molecular mechanisms resulting in dementia(Kayed & Lasagna-Reeves, 2013). The oligomers have a number of neurobiological targets, and prominent among them are α7 nicotinic acetylcholine receptors(Caterina M. Hernandez et al., 2010; Inestrosa et al., 2013; Lilja et al., 2011; Sadigh-Eteghad et al., 2014). We hypothesize that the interaction of beta amyloid products with α7 nicotinic receptors is central to some pathologies of Alzheimer’s dementia.

Our recent nonhuman primate work demonstrated that long-term activation of the cholinergic basal forebrain resulted in cognitive improvements(Blake et al., 2017a; Liu et al., 2017, 2018; Qi et al., 2021). One key observation was that one hour of intermittent stimulation of the cholinergic basal forebrain, for multiple days of the week over months, led to changes in working memory duration that were adequate to change a monkey’s rank percentile by 32-44% (Blake et al., 2017b; Liu et al., 2017). Although these effects were reliably induced and substantial, mechanisms of change over those time scales from cholinergic modulation were unclear.

We set out to test hypotheses on mechanisms of these improvements by performing long-term stimulation in mice. We chose mice to facilitate protein expression analysis. We chose the 5xFAD mouse for the Alzheimer’s model because it aggressively deposits Aβ_42_ and has a genotype-dependent decline in cognition (Oblak et al., 2021). By applying deep brain stimulation of the cholinergic basal forebrain, we hoped to see behavioral improvements dependent on stimulation, and to be able to investigate protein expression key to the cholinergic signaling and beta amyloid metabolic pathways.

## MATERIALS AND METHODS

### Test Subjects

B6SJL-Tg(APPSwFlLon,PSEN1*M146L*L286V)6799Vas/Mmjax or 5XFAD were used for study. colony. Transgenic 5XFAD and wildtype littermates were identified by PCR using a sense primer (5’- CTA CAG CCC CTC TCC AAG GTT TAT AG-3’) and antisense primers (5’- AAG CTA GCT GCA GTA ACG CCA TTT-3’) and (5’- ACC TGC ATG TGA ACC CAG TAT TCT ATC-3’). PCR products were separated on a 2% agarose gel, and a DNA ladder confirmed correct band size. Hemizygous mice of both genotypes and 5XFAD non-transgenic control littermates were housed for four months before all experiments. Mice were housed in groups of 5 per cage, in a temperature and humidity controlled room on a 12-hour light and dark cycle (06:00-18:00). Animals were given free access to food (Teklad Rodent Diet 8604 pellets, Harlan, Madison, WI) and water. All testing was performed during the light cycle portion of their day. The Institutional Animal Care and Use Committee of Augusta University approved of all protocols and procedures used in this study. Under the guidance of the 8^th^ Edition of the National Institute of Health Guide for Care and Use of Laboratory Animals (2011), measures were taken to minimize pain and discomfort throughout the study.

### Electrode Fabrication

Stimulation electrodes were custom-made based on previously published specifications(McCairn & Turner, 2009). Stainless steel 30 ga. hypodermic tubing (MicroGroup ®, Medway, MA) was cut to a length of 2.8mm using a Dremel ® rotary saw. A 26G (Precision Glide ®) needle was twisted into each end of the tubing to re-establish their patency and a tungsten wire reamed out any other debris. Teflon coated Platinum-iridium wire with an outer diameter of 139.7um (A-M Systems, Seattle, WA) was cut to 20mm in length and threaded through the hypodermic tubing. Iris scissors grasped and unsheathed 1.0mm of the Pt-ir wire to create the conductive tip. Polyimide tubing was cut to a 3.8mm length and slide onto the hypodermic tubing and wire so that the 1.0mm excess polyimide tubing was left at the tail end of the electrode. The stimulation tip was adjusted in the hypodermic tubing exposing 0.5mm of uncoated wire. Using the included brush, cyanoacrylate (Krazy ®) was applied to the excess polyimide opening until glue was drawn into the area creating a 4.8mm long electrode from the stimulation tip to end of polyimide tubing. A butane torch uncoated about 3mm of the tail-end wire. Electrodes were left to dry overnight.

### Surgical Procedures

#### Stimulation Electrode and Headcap Implantation

Surgery was performed on a sterile field under 1.5% isoflurane anesthesia. Artificial tears ophthalmic ointment (Akorn, Lake Forest, IL) was applied to both eyes. After shaving the scalp with hair clippers, the mouse was placed into a bell jar with 0.5mL isoflurane until loss of righting reflex. The mouse was transferred to a dual arm, rodent stereotaxic instrument (Stoelting ®, Wood Dale, IL) fitted with an anesthesia mask and heating pad set to 37°C. A 10% povidone-iodine Betadine Swabstick (Purdue ® Stamford, CT) was applied to the scalp then cleaned off with 70% isopropyl alcohol prep pads. Surgical scissors removed the scalp, and a periosteal elevator scraped clean the periosteum. Stereotaxic measurements were made in reference to the bregma landmark (−0.6mm AP or caudal, 1.7mm ML or lateral) to target sites for electrode implantation. A 0.5 mm hole was drilled at target sites using a carbide bur (HM1-005-FG, Meisinger, Centennial, CO) and dental drill (Volvere GX, Brasseler, Savannah, GA) until the cranium was breached. An electrode was loaded into a guide tube positioned above the target site and pushed into the cerebrum with a stylus until the back of the electrode became flush with the cortical surface to ensure achievement of 4.8mm depth. After raising the stylus and guide tube, cyanoacrylate was placed at the target site to stabilize the electrode and seal off exposed tissue. The procedure was repeated for implantation of the second electrode. With the same burr, a hole was made at each temporal and frontal bone and stainless steel screws (Antrin, Fallbrook, CA) were hand screwed two rotations. Before implanting the final bone screw in the left frontal bone, a 20mm length of pt-ir wire with 3mm of coating removed at each end was procured. One end was bent with forceps and then slid into the bone screw hole so that the tip resided between the top of the cortex and bottom of the skull. The bone screw was then placed as usual. A layer of cyanoacrylate was applied to all screws and electrode sites and left until dry. Methyl methacrylate powder (A-M Systems, Everett, WA) was mixed with its respective dental cement solvent until saturated and then a thin layer of the mix was applied to the top of the skull with the periosteal elevators. A type B micro-USB receptacle (part no. 84×0185, Newark®, Chicago, IL), with the 2^nd^ and 4^th^ connector pins removed, was fixed to one of the stereotaxic arms and positioned over the skull. The exposed wire from the bone screw was tied to the grounding pin of the micro-USB port while the electrode wires were tied to the remaining two pins. Each wire was connected to its pin with a soldering iron. A thin coat of dental cement was applied to the soldered pins to ensure firm connection. The micro-USB receptacle was dismounted from the stereotactic arm and positioned just above the top of the bone screws. A larger stainless steel screw (MX-M025, Small Parts Inc., Logansport, IN) was adhered to the side of the micro-USB receptacle that would serve as a handle when connecting mice to the stimulating computer. Dental cement was applied to fill in all areas between the base of the skull and base of the micro-USB receptacle to create the stimulating headcap. A micro-USB receptacle was placed into the top of the headcap to prevent cage-mates from chewing on the internal connector. The mouse was returned to the cage and monitored until conscious and consuming water and food. Control mice underwent identical procedures, however the electrode and grounding wires were clipped instead of attaching to the micro-USB receptacle pins.

Lesions were created in pilot animals developing implantation approaches until targeting was consistent. Lesions were created with a 30 μA DC current, electrode negative, for 10 seconds which generated a clearly visible mark on perfused and unstained sections.

#### Viral Transduction

Surgery was performed on a sterile table under 1.5% isoflurane anesthesia. Artificial tears ophthalmic ointment was applied to both eyes. A shaved mouse was transferred to the rodent stereotaxic instrument. The scalp was sanitized as previously mentioned and then a 3mm incision was made with surgical scissors at an approximated location to expose the skull around 2mm lateral and posterior to bregma. A small hole was carefully made (−2.0mm AP, 2.0 mm ML from bregma) over the somatosensory cortex so that no bleeding events occurred which would interfere with adeno-associated virus delivery. A micropipette was slowly inserted over 5 minutes to a depth of 700μm. Non-transgenic 5XFAD littermates were injected with 400nL of (AAV9-hsyn-ACh4.3, Vigene) at 100nL/min using a micropump (WPI). After a 5-min wait period, the micropipette was slowly withdrawn from the cortex over 5 minutes. The scalp incision was sutured closed and sealed using tissue adhesive (Vetbond™, 3M). Mice were returned to the home cage, and the viral vector incubated for 3-wks. To confirm vector expression, mice were anesthetized and prepped as stated above. The scalp was re-incised to expose the injection sight, and the presence of fluorescence was checked using confocal epifluorescence microscopy fitted with a FITC filter (emission of 513-556nm, excitation of 467-498nm). Mice demonstrating positive fluorescence and thus GACh expression were then implanted with an optical window and single microelectrode respectively detailed below.

#### Optical Window and Electrode Implantation

Implantation of cranial window was performed as described previously (Mostany et al., 2008). Briefly, after mice were anesthetized with isoflurane (4% induction, 1.2-1.5% maintenance), mice were shaved and then skin and connective tissue attached to the skull were removed. To form an ∼3 × 3 mm round cranial window, cranial bone over the somatosensory cortex was carefully removed using a ¼ round carbide dental bur (Midwest). The brain was then irrigated with sterile cortex buffer containing the following (in mM: 135 NaCl, 5.4 KCl, 1 MgCl2, 1.8 CaCl2, and 10 HEPES, pH 7.3), and a 5-mm-diameter #1 glass coverslip (Electron Microscopy Sciences) was placed over the window and sealed to a custom-made metal ring using cyanoacrylate and dental cement. A single 6.8 mm microelectrode was fabricated and inserted as described previously but at -5.4 mm AP and 1.7 mm ML to bregma and implanted at a 45° angle to the horizontal plane in order to stimulate the nbM without obstructing the optical window. The head ring was fit into a custom-made stereotaxic holder for imaging under isoflurane anesthesia.

#### Two Photon Imaging

The Nikon A1R MP multiphoton system mounted on the FN1 upright microscope was used to collect images with a 25x/1.1 NA water-immersion objective. The Spectra-Physics Mai-Tai EHP laser tuned to 950 nm was used for two-photon excitation and emission light was collected by a GaAsP detector using a bandpass filter (500–550 nm). The time-lapse images consisting of a single section were taken at 1 frame per second for 5 minutes across 78 × 78μm imaging field within layer I of the sensorimotor cortex.

#### Pericardial Perfusion

Mice are placed in a bell jar with 0.5 mL of isoflurane until loss of righting reflex and then transferred to an isoflurane vaporizer at 2% concentration. After a 10-min period to achieve a sufficient plane of anesthesia and unresponsive toe-pinch, mice are placed supine onto a Styrofoam surface. A 20G (Precision Glide ®) needle was inserted through each extremity to mount the mouse to the surface. A 6 cm lateral incision was made through the integument and abdominal wall inferior to the ribcage. An incision is made in the diaphragm and curved scissors make incisions through both sides of the ribcage to the clavicle bones. The sternum and ribs are reflected away from the chest cavity and blunt dissection exposed the heart. Iris scissors made a small incision in the right atrium to allow outflow of systemic blood. The 26G needle attached to the perfusion apparatus was inserted into the left ventricle and a pressure of 115 mmHg was maintained as 50 mL of 4% paraformaldehyde solution at 4°C was perfused throughout the body.

### Brain Stimulation

#### Electrode Impedance Measurement

The individual impedances of each electrode had to be calculated so that the stimulation voltages could be tailored to deliver 100 μA of current to each electrode tip. Because previous pilot studies demonstrated a consistent daily impedance, impedances were measured at the beginning of each week to continue to ensure their long-term consistency. Impedance was measured using custom programmed software that triggered a stimulus isolator (A385 World Precision Instruments, Sarasota, FL) to produce a known 100 μA current square wave and then use a connected oscilloscope (TDS 2012C Tektronix, Beaverton, OR) to measure the voltage change. After confirming calibration of the method using a known impedance of a circuit resistor, the same method was applied to the left and right electrodes of each mouse head cap. Impedances averaged 8-14 kOhm. Variable resistors were added to the stimulation cord circuit path when needed in order to stimulate multiple mouse electrodes at the same time with a uniform voltage output. Typically, stimulation was delivered at close to 1V peak per phase across an impedance of 10 kΩ.

#### Mouse Stimulation

Mice were brought to the stimulation room and allowed 30-min of acclimation before beginning stimulation. Remaining in their home cages, the large screw adhered to the side of the micro-USB receptacle was secured with hemostats while a micro-USB cable was plugged into each mouse headcap. The mouse cage lids were returned ajar so that mice had continued access to food and water. All micro-USB charging cables were threaded through a pulley system equipped with a counter balance and suspended from the ceiling to remove any upward or downward forces on the headcaps during stimulation and allowing for better mouse mobility. Tantalum capacitors (10μF 25v, Kemet, Fort Lauderdale, FL) were soldered to each stimulating cable end to block net charge transfer. These capacitors and the cable’s grounding wires were connected to a multiple functional I/O device (USB-6211, National Instruments, Austin, TX) that served as a voltage generator creating the voltage-dependent stimulation pulses signaled by custom programmed software. Stimulation was delivered with biphasic, negative first, unipolar 100 μA pulses with 100 μS per phase. 60 pulses were delivered per second for 20-sec followed by 40-sec of no stimulation. Mice received 1-hr of bilateral stimulation 5-days each week.

### Tissue Processing

#### Brain Extraction

Mice were placed in a bell jar with 0.5 mL of isoflurane until loss of righting reflex and unresponsive to toe pinch. Grasping the base of the skull and tail, manual cervical dislocation was performed to euthanize the mouse. Surgical scissors transected through cervical tissue and spinal cord in order to isolate the head. The scalp was cut and pulled away exposing bare cranium. Incisions were made laterally at the foramen magnum and continued transversally around both sides of the cranium. The headcap, attached to the temporal and frontal bones, was pulled upwards exposing the brain. A periosteal elevator removed the brain from the cranium and transferred it face down onto a sterile petri dish on a bed of crushed ice. A #11 scalpel made a coronal incision at the origin of the cranial nerve II white matter tracts. The cortical portion of the anterior section was removed with forceps and a scalpel. The cortical portion of the posterior brain section was reflected from the midline using a blunt probe to expose both hemispheres of the hippocampus which were resected with forceps. All frontal cortex and hippocampus tissue was separated by hemisphere and transferred to labeled 1.7 mL microcentrifuge tubes on ice and then stored at -80°C.

#### Tissue Homogenization

100 μl of 4°C radioimmunoprecipitation buffer (Sigma-Aldrich, St. Louis, MO) with 10 μl/ml protease inhibitor cocktail (AEBSF at 104 mM, Aprotinin at 80 μM, Bestatin at 4 mM, E-64 at 1.4 mM, Leupeptin at 2 mM and Pepstatin A at 1.5 mM, Sigma-Aldrich, P8340) and was added to each frozen brain section on ice. Samples received a 5-sec pulse from an ultrasonic homogenizer (Qsonica ®, Newtown, CT) while sitting in crushed ice. Homogenized samples underwent centrifugation at 15,000 g for 15-min at 4°C. Supernatants were transferred to pre-chilled microcentrifuge tubes and stored at -80°C until used for protein analysis.

### Protein Analysis

#### Western Blot Analysis

Protein concentrations of each sample were determined using the bicinchoninic acid (BCA) protein assay kit (Thermo Fisher Scientific, Waltham, MA) which used bovine serum albumin standards to create a best fit linear regression line based on protein concentrations using the 562nm absorbance mode of a microplate reader and its Gen5 Data Analysis program (Biotek, Winooski, VT). Equal amounts of protein were resolved in SDS-polyacrylamide gels and electrophoretically transferred to nitrocellulose membranes (Bio-Rad, Hercules, CA) at 4°C. Membranes were blocked for 1-hr in Tris-buffered saline containing Tween 20 (TBST; 10mM Tris-HCl, pH 8.0, 138 mM NaCl, 2.7 mM KCl, and 0.05% Tween 20) and 5% non-fat milk.

Primary antibodies used were anti-BDNF (1:1000, GTX134514, GeneTex ®, Irvine, CA), anti-NGF, (1:1000, AN-240, Alomone Labs), anti-TrkA (1:1000, GTX132966, GeneTex ®, Irvine, CA), anti-pTrkA-Y490 (1:1000, product no. 9141S, Cell Signaling Technology ®, Danvers MA), anti-TrkB (1:1000, GTX54857, GeneTex ®, Irvine, CA), anti-pTrkB-Y705 (1:1000, ab229908, Abcam, Cambridge, MA), anti-BACE1 (1:1000, GTX103757, GeneTex ®, Irvine, CA), and anti-ADAM10 (1:1000, GTX104940, GeneTex ®, Irvine, CA).

Primary antibodies were added to 10 ml of 5% non-fat milk and TBST solution and applied to the blocked membrane to incubate overnight on an orbital shaker at 4°C. The following day, membranes were washed three times with TBST for 5-min and then incubated for 1-hr with horseradish peroxidase-conjugated goat anti-rabbit IgG (ab6721, Abcam, Cambridge, MA). Membranes were washed three more times with TBST for 5-min.

Blots were developed using Pierce™ ECL Western Blotting Substrate (ThermoFisher, USA, Cat# 32106) at a sufficient volume to ensure that the blot was completely wet with the substrate and the blot didn’t become dry (0.1 ml/cm^2^). Membrane incubated with the substrate working solution for 1 minute. Blot was removed from ECL solution and placed in a plastic sheet protector or clear plastic wrap. An absorbent tissue removed excess liquid and carefully pressed out any bubbles from between the blot and the membrane protector. Then the blot was analyzed using The ChemiDoc XRS+ Gel Imaging System (BioRad, USA). The images were analyzed using Image Lab image acquisition and analysis software (BioRad, USA).

To avoid saturation and ensure linearity, 30μg of each sample were loaded in the gel. Study has shown that GAPDH showed linearity up to 30 μg of protein loaded. Along with this, gels were stained with Coomassie stain to ensure equal amount of protein loaded (PMID: 24023619, 26657753).

For the different molecular weights, membranes were cut into strips (horizontally using protein ladder/marker), then probed individually with different antibodies. However, for very close molecular weights and/or for the GAPDH, membrane blots were stripped using western blot stripping buffer (Thermo Fisher, USA, cat# 21059). For stripping, blots were washed 5 min x 3 with TBST to remove chemiluminescent substrate, and then incubated in western blot stripping buffer for 5 to 15 minutes at RT. Then they were again washed 5 min x 3 with TBST, and then performed next immunoblot experiment.

#### Enzyme-linked Immunosorbent Assay

Following the manufacturer’s instructions for the mouse Aβ_42_ ELISA kit (Invitrogen ®, Waltham, MA), a solution of guanidine-HCl in 50mM Tris (pH 8.0) was added to homogenized brain samples to achieve 5 M guanidine-HCl and mixed for 4-hr on orbital shaker at room temperature. Samples were quantified with the BCA kit and diluted with the included diluent buffer to achieve kit recommended concentrations. Samples were then added to the well plate, coated with monoclonal antibody against the NH2 terminus of Aβ_42_, covered with an adhesive plate cover, and incubated for 2-hr at room temperature. 100μl of Ms Detection Antibody solution was added to each well. The plate was re-covered and incubated at room temperature for 1-hr. Wells were washed four times using a squeeze-type wash bottle with the provided Wash Buffer. 100μl of Anti-Rabbit IgG HRP solution was added to each well and incubated for 30-min. Wells were rewashed as previously stated. 100μl of Stabilized Chromogen was added to each well and incubated for 30-min before adding 100μl of the provided Stop Solution. Aβ_42_ standards were included in this protocol to create a best-fit standard curve and sample absorbance was detected at 450 nm using the microplate reader. Aβ_42_ sample concentrations were calculated by dividing the Aβ_42_ concentration from the ELISA kit by the total protein concentration from the BCA kit.

### Water Maze

#### Test Apparatus

The water maze consisted of a circular tank (1.80-m diameter and 0.76-m height) made of black plastic and filled to a depth of 35 cm of water. An escape platform (20-cm diameter) was placed 1 cm beneath the surface of the water and centered in one of the four tank quadrants. Water (maintained at 22°C ± 1.0 °C) was made opaque with nontoxic white paint. The maze was surrounded by visual cues including a green watering can, blue lab coat hanging from a standing hanger, and black curtains, on which white paper geometric images were affixed to provide visual cues. These curtains also hid the experimenter and resting test subjects from view. Swimming activity was monitored using an overhead mounted camera that relayed mouse swimming location and trajectory as well as latency to platform measurements to a video tracking system (Noldus EthoVision ® Pro 3.1).

#### Hidden Platform Task

For each test day, mice were transferred to new testing cages and then given 30 minutes to acclimate to the water maze room surroundings before starting the first trial of the day. For each daily test session, mice were given 90-seconds per trial to locate and climb onto the escape platform. A trial was initiated by placing each mouse into the water directly facing the center of the maze pool at one of 7 locations around the pool wall. Starting locations were pseudorandomized so that starting locations were equally distributed among the four quadrants for every mouse over the 5-day testing period. Mice that found the platform successfully were given a 30-second rest period on the platform before being returned to their testing cage. Mice that did not find the platform within the 90-second trial were guided to it using a curved plastic handle and then given a 30-second rest period before returning to the testing cage. Mice were given a 30-minute rest in their training cage before undergoing the subsequent trial. Each day consisted of 3 trials per day for five consecutive days.

#### Probe Trials

After 24 hours from the last hidden platform task, mice were returned to the water maze room with a 30-minute acclimation period. The hidden platform was removed from its quadrant in the pool. For this single trial, each mouse was placed at the same start position in the quadrant opposite of the previously hidden platform quadrant. For the observer on the tracking monitor, the video tracking system projected a 30 cm radius circle over the previous location of the escape platform. Each mouse was given 90 seconds in the pool, and the observer counted how many times each mouse enters and then re-enters the probe test zone.

#### Visible Platform Task

To confirm general visual acuity in test animals, a screwdriver was placed face down through one of the escape platform holes so that the red handle was raised 12 cm above the water surface. After a 30-min period after completing the probe trial, each mouse was placed in the same starting location and time to swim to the platform was measured.

### Accelerating Rotarod

#### Test Apparatus

Motor coordination and balance were tested using an accelerating rotarod (Rotor-Rod System ®, San Diego Instruments).

#### Rotarod Training

Mice were placed into testing cages and then brought to the testing room to acclimate for 30-min. Each mouse was then placed on the station rod for 10-sec before starting the rotation motor. The rod was then gradually increased from 0-10 rpm over a 1-min period and maintained at 10 rpm for 5-min. During the first trial, mice that fell off of the rod were placed back onto it until the end of the 5-min duration. Mice performed three trials total with an intertrial interval of 20-min and latency to fall was recorded for each. Trials where mice hang onto the rod and make two continuous rotations without any further footsteps would be considered a fall and the latency also recorded. Mice were then returned to their home cages and room.

#### Rotarod Test

After 24-hrs from training, mice were placed into training cages and brought to the rotarod room for a 30-min acclimation period. Each mouse was given a single training trial as described above in order to refresh them of the task and returned to their testing cages. After a 20-min rest, mice were returned to the rod which would now increase from 0-23 rpm over a 2-min period during the 5-min trial. Mice underwent three trials with 20-min rest in between each trial and latency to fall was recorded for each mouse. Trials where mice made two consecutive rotations without taking a step were recorded as a fall.

### Statistical Analysis

Statistical analyses were performed using GraphPad Prism 9 software.

## Results

### Acetylcholine release documented with stimulation

Groups of wild-type mice had the vector GACh4.3 infused into the cerebral cortex. After verification of expression, animals were implanted with an optical window, and mesoscale fluorescent imaging was conducted. Expression of the receptor is shown in Figure 1A. Test stimulation was conducted to verify responsiveness to pulse trains. Responses to pulses from three animals are shown in Figure 1D-F. In two of the animals, a comparison of continuous 60 pulse per second stimulation was performed, and compared to intermittent stimulation at the same pulse rate, cycled on for 20 seconds and off for 40 seconds of each minute. The onset of stimulation, or onset of a minute of intermittent stimulation, generally led to 10% increases in fluorescent signal. In the two animals in which continuous stimulation was tested, mesoscale fluorescence returned to baseline within minutes, while intermittent stimulation elicited a phasic response to each 20 second on-cycle. A color coding of the mesoscale image was performed during one example of the intermittent stimulation, and the differences between the end of the off-cycle, and the peak of the on-cycle, are shown in Figure 1B-C. These evidence contributed to confidence that the stereotaxic location chosen and verified with lesions was appropriate to induce acetylcholine release in vivo.

**Figure 1.**
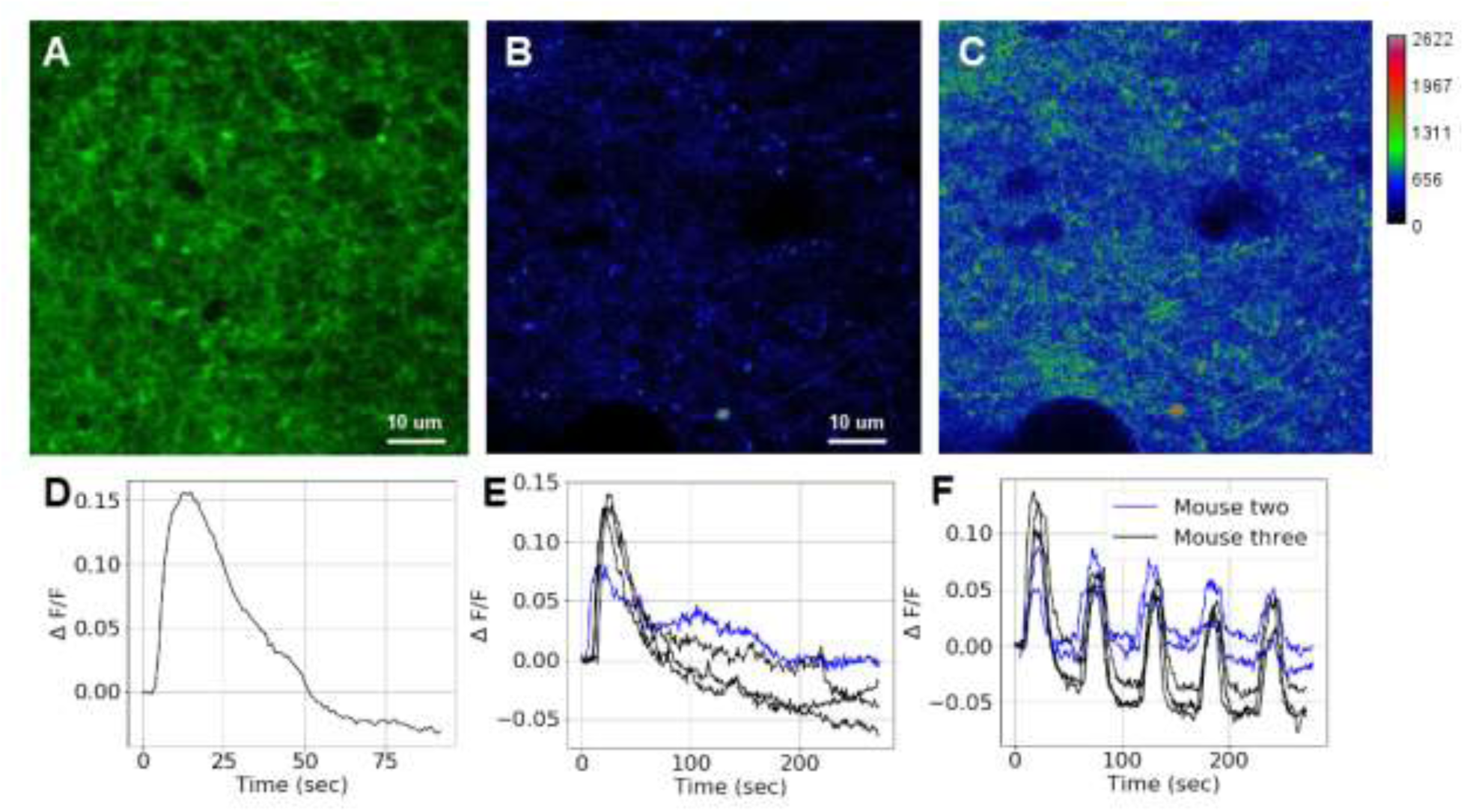
GACh4.3 responses. A. Cortical expression of the GACH4.3 reporter focused 20 microns below the surface. B. Single frame color-coded fluorescence intensity from a sample image at the nadir of intensity in F. C. As in B), color-coded fluorescence intensity at the peak. D. Phasic response to 20 seconds of pulse stimulation in mouse one. E. Overlaid mesoscale average-pixel fluorescence intensity traces from mouse two and three showing responses to continuous 60 pulse per second stimulation. F. As in E, for 60 pulse per second stimulation that cycles on for 20 seconds and off for 40 seconds. E and F show single trial mesoscale data.

### Behavioral results

Groups of wild-type and 5xFAD mice were subjected to stimulating electrode implantation of the cholinergic basal forebrain. A series of lesions were performed in pilot animals until macrohistological reconstructions demonstrated consistent targeting, as shown in Figure 2A. Animals were tested in the Morris Water Maze(Morris, 1984) using three randomized starting points each day, with a 90 second timeout. Swim times are displayed in Figure 2B. A three way ANOVA found significant effects of genotype (F(1,115)=11.06, P=0.0012), day (F(4,115)=18.5, p<0.0001), and stimulation (F(1,85) = 35.35, *p* < 0.0001). A posthoc Fishers test found that the wild-type stimulated group had lower swim times on days 4 and 5 (wt *t*(120) = 3.21, *p* < 0.0086; *t*(120) = 2.94, *p* < 0.0199) than the wild-type unstimulated, and in the 5xFAD groups on days 3-5(*t*(120) = 3.38, *p* < 0.0057; *t*(120) = 3.51, *p* < 0.0038; *t*(120) = 3.62, *p* < 0.0027). On the water maze probe test (Figure 2E), the stimulated 5xFAD group crossed the platform location significantly more often than unstimulated groups (wt t=4.13,p<0.002, 5xFAD t=4.07,p<0.002).

**Figure 2.**
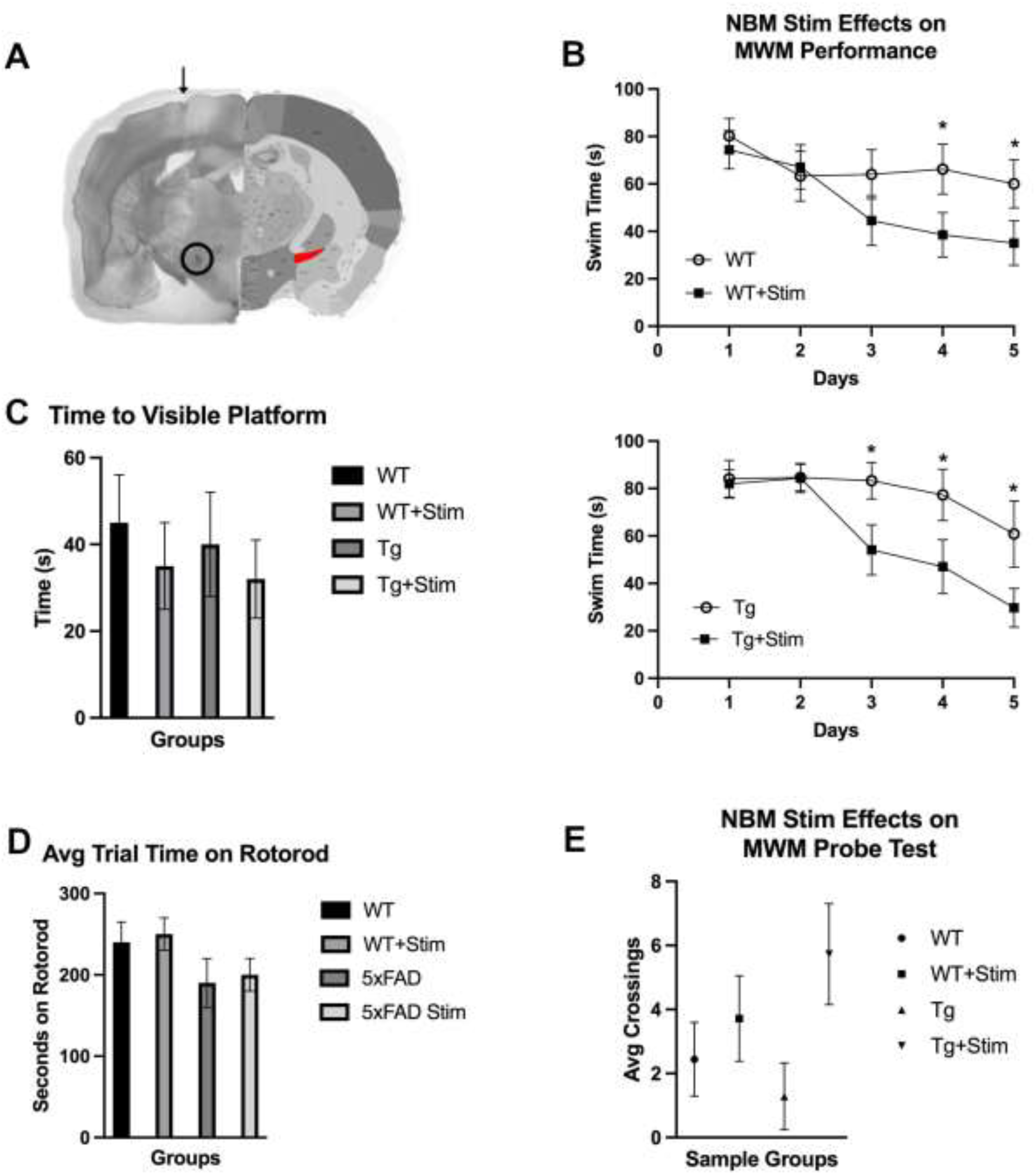
(A) Electrolytic lesion mapping. Coronal wet specimen of mouse brain after delivery of anodal current pulses to induce an electrolytic lesion at site of exposed microelectrode tip implanted -0.6 mm AP, 1.7 mm ML, and 4.8 mm DV to bregma. Black arrow: cortical site of microelectrode penetration. Black circle: electrolytic lesion. Red region: substantia innominata region of intended stimulation from Allen Mouse Brain Reference Atlas. (B) Effects of nbM stimulation on water maze escape latency. 3-way ANOVA determined significant difference in average escape latency for the individual effects of days *F*(4,115) = 18.49, *p* < 0.0001, genotype *F*(1,115) = 11.06, *p* < 0.0013, stimulation *F*(1,85) = 35.35, *p* < 0.0001, and a significant between-factor effect of days x stimulation *F*(4,85) = 5.33, *p* < 0.0008. Fisher’s LSD *post hoc* multiple comparisons with a Bonferroni correction determined stimulation significantly decreased mean escape latency during water maze acquisition on days 4-5 for wild-type mice *t*(120) = 3.21, *p* < 0.0086; *t*(120) = 2.94, *p* < 0.0199 respectively (upper panel), and on days 3-5 for 5xFAD mice *t*(120) = 3.38, *p* < 0.0057; *t*(120) = 3.51, *p* < 0.0038; *t*(120) = 3.62, *p* < 0.0027 respectively (lower panel). (C) Visible platform test. Mean escape latency to visible platform for nbM stimulated and unstimulated 5xFAD and wild-type mouse groups. One-way ANOVA determined no significant difference between groups *F*(40) = 0.32, *p* < 0.81. (D) Accelerating Rotarod performance. Mean time spent balancing on rotating rod for each mouse group. One-way ANOVA determined no significant difference between groups *F*(40) = 1.40, *p* < 0.26. (E) Water maze probe test. Mean number of mouse crossings over the removed escape platform per mouse group. One-way ANOVA followed by a *post hoc* Fisher LSD test with Bonferroni multiple comparisons correction determined that nbM stimulation caused a significant increase in platform crossings in the 5xFAD group *t*(40) = 4.13, *p* < 0.002. Control wild-type mice: n = 6 males & 6 females. Stimulated wild-type mice: 7 males and 7 females. Control 5xFAD mice: n = 3 males and 4 females. Stimulated 5xFAD mice: n = 9 males and 2 females. * and ** indicate *p* < 0.05 and 0.01 respectively.

Control behaviors were used to assess the likelihood of differences based in motor ability rather than memory. Visible platform swim results are presented in Figure 2C. A one way ANOVA did not find significant effects of stimulation or genotype on swim times (*F*(40) = 0.32, *p* < 0.81). Time to fall on the rotorod was tested, and results are shown in Figure 2D. A one way ANOVA did not find significant effects of group (*F*(40) = 1.40, *p* < 0.26).

A ten minute alternating Y-maze test was performed. Two way ANOVAs on percentage of alternation and number of arm entries did not show statistical differences based on stimulation treatment (F(1,35)=0.25 p=0.62 and F(1,35)=0.55 p=0.46 respectively). Transgenic animals made fewer arm entries (F(1,35)=7.6, p<0.0009).

### Nerve Growth Factor and related receptors

The release of acetylcholine in the cortical mantle has been reported to increase nerve growth factor signaling through its activation of nicotinic acetylcholine receptors (Hotta et al., 2009; Jonnala et al., 2002). To assess whether these pathways were altered by the five months of stimulation, animals were left unstimulated for over 24 hours prior to tissue harvest to allow short term activation related processing to finish(Kaplan et al., 1991). Frontal cortex was probed using antibodies for mature nerve growth factor (mNGF), precursor NGF (proNGF), the tropomyosin receptor kinases receptor (TrkA), and the activated TrkA receptor as indicated by its phosphorylation (pTrkA).

Figure 3A-C shows the changes in mNGF and proNGF. The mature form increased in both groups of stimulated animals compared to unstimulated littermates of the same genotype, and this effect reached significance in the wild-type animals (wt t(4) = 9.15, p < 0.0001; 5xFAD t(4) = 1.28, p < 0.271). Figure 4D shows significant increases in TrkA in both groups of stimulated animals compared to unstimulated controls of the same genotype (wt t(4) = 2.96; p < 0.042. 5xFAD t(4) = 4.74; p < 0.01). Figure 4E shows increases in pTrkA caused by stimulation in both genotypes, and this effect reached statistical significance in the wild-type groups(wt t(4) = 2.83; p = 0.048; 5xFAD t(4) = 0.622, p < 0.57).

**Figure 3.**
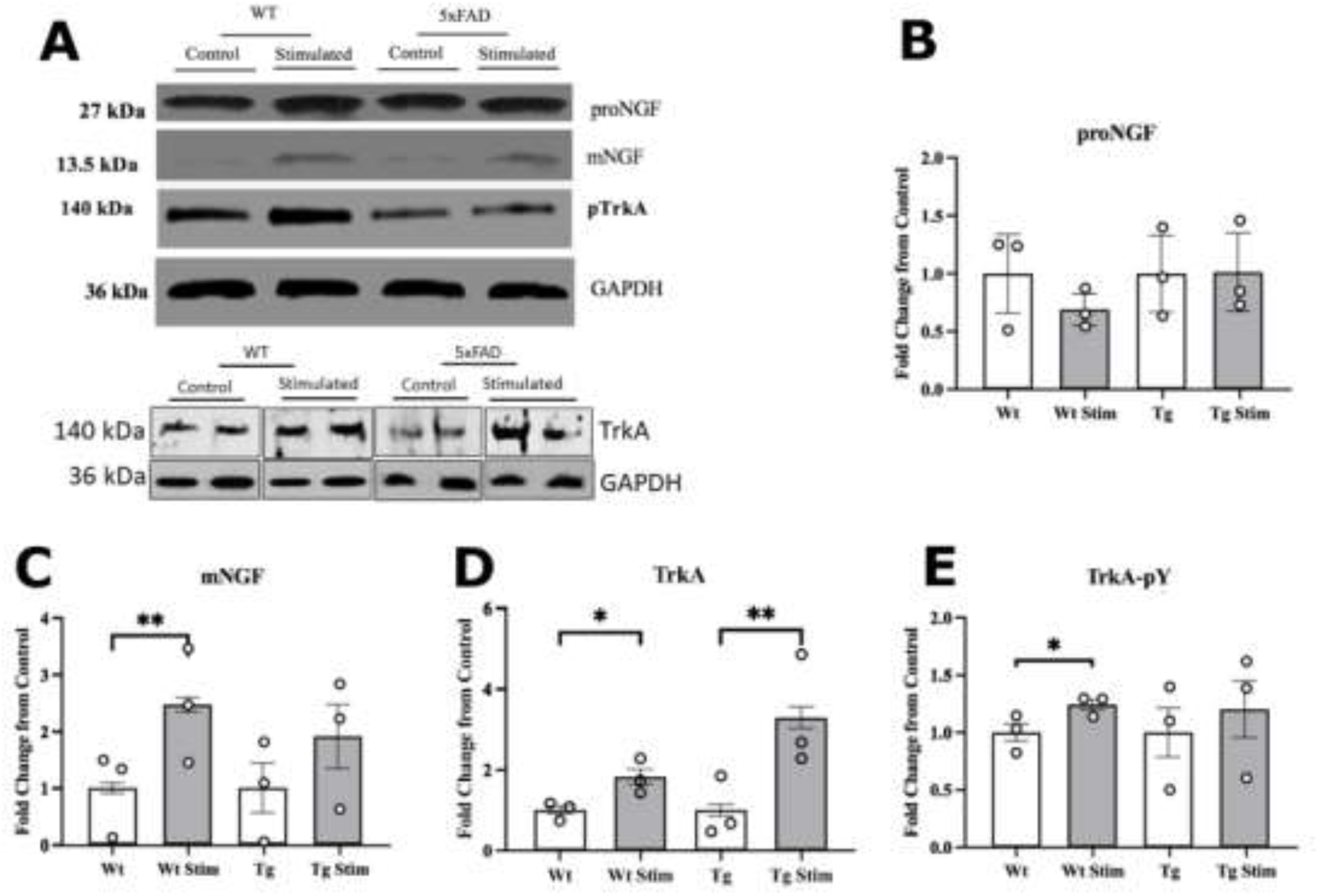
Concentrations of nerve growth factor pathway markers after nbM stimulation. Frontal cortex tissue was homogenized from of WT and 5xFAD mice and immunoblotted using antibodies described in the Materials and Methods section. (A) Representative immunoblots. (B-E) Optical density values normalized against GAPDH and expressed as a percentage of the control group. Data is represented as mean per group ± SEM. *n* = 3 animals per group. (B) Immunoblot of pro-NGF at 27 kDa. The WT group showed a non-significant decrease in proNGF. WT t(4) = 0.854, p < 0.442; 5xFAD t(4) = 0.026, p < 0.99. (C) Immunoblot of mNGF at 13.5 kDa. Both groups showed increase in mNGF with the WT group reaching significance. WT t(4) = 9.15, p < 0.0001; 5xFAD t(4) = 1.28, p < 0.271. (D) Immunoblot of NGF’s receptor TrkA. Both the WT and 5xFAD groups showed significant increases. WT t(4) = 2.96; p < 0.042. 5xFAD t(4) = 4.74; p < 0.01. (E) Immunoblot of the phosphorylated active form of TrkA. Both groups showed increase in the phosphorylated TrkA with significance reached in the WT group. WT t(4) = 2.83; p = 0.048; 5xFAD t(4) = 0.622, p < 0.57. * and ** represent p < 0.05 and p < 0.01 respectively.

**Figure 4.**
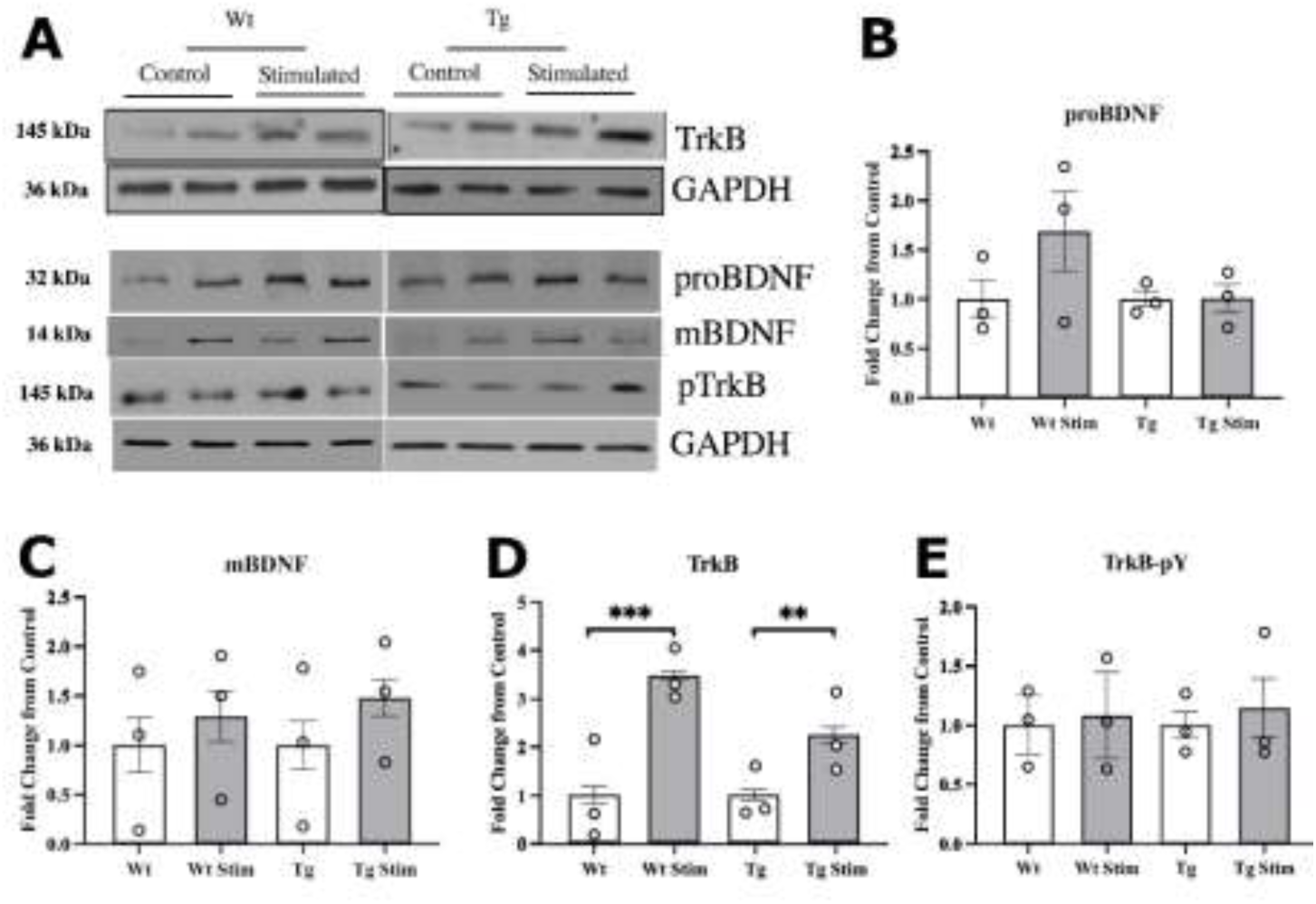
Concentrations of the brain derived neurotrophic factor pathway markers after nbM stimulation. Proteins were immunoprecipitated from frontal cortex of WT and 5xFAD mice and immunoblotted using antibodies described in the Materials and Methods section. (A) Representative immunoblots. (B-E) Optical density values normalized against GAPDH and expressed as a percentage of the control group. Data is represented as mean per group ± SEM. *n* = 3 animals per group. (B) proBDNF. WT t(4)=1.54, p<0.20 5xFAD t(4)=0.049, p<0.96 (C) mBDNF. WT t(4)=0.77 p<0.49 5xFAD t(4)=1.58 t<0.19 (D) TrkB at 140 kDa. Both groups showed a significant increase in TrkB. WT t(4) = 11.94, p < 0.0003***. 5xFAD t(4) = 6.21, p < 0.003**. (E) pTrkB WT t(4)=0.181 p<0.86 5xFAD t(4)=0.54 p<0.62 * and ** represent p < 0.01 and p < 0.001 respectively.

### Brain derived neurotropic factor and related receptors

The brains of Alzheimer’s patients have lower levels of TrkB than age matched controls (Allen et al., 1999). We queried whether frontal cortex TrkB protein levels were altered by basal forebrain stimulation. Figure 4D shows that stimulation caused significant two to four-fold increases compared to unstimulated littermates of the same genotype, and this effect was significant in wild-type and 5xFAD genotypes(wt t(4) = 11.94, p < 0.0003. 5xFAD t(4) = 6.21). Nonsignificant results were obtained for the precursor brain derived neurotrophic factor (proBDNF), mature brain derived neurotrophic factor (mBDNF), and the phosphorylated TrkB receptor as shown in Figure 4.

### Amyloid processing pathway measures

The transgenic genotype used was the 5xFAD genotype(Oakley et al., 2006), which has three amyloid precursor protein mutations and two presenilin mutations to result in beta amyloid production and accumulation. To assess the impact of stimulation on the presence of amyloid pathway elements, we probed frontal cortex tissue for the proteins ADAM10, BACE1, and for beta-amyloid. Western blotting for ADAM10, the alpha cleavage enzyme, shown in Figure 5A did not find significant changes caused by stimulation. BACE1, the beta-amyloid cleavage enzyme, trended down in both groups but was not significant at the 0.05 level as shown in Figure 5B. An ELISA measured concentration of Aβ_42_ found levels in the tissue were reduced significantly, and almost 50% (5xFAD t(6) = 12.77, p < 0.001; Figure 5C).

**Figure 5.**
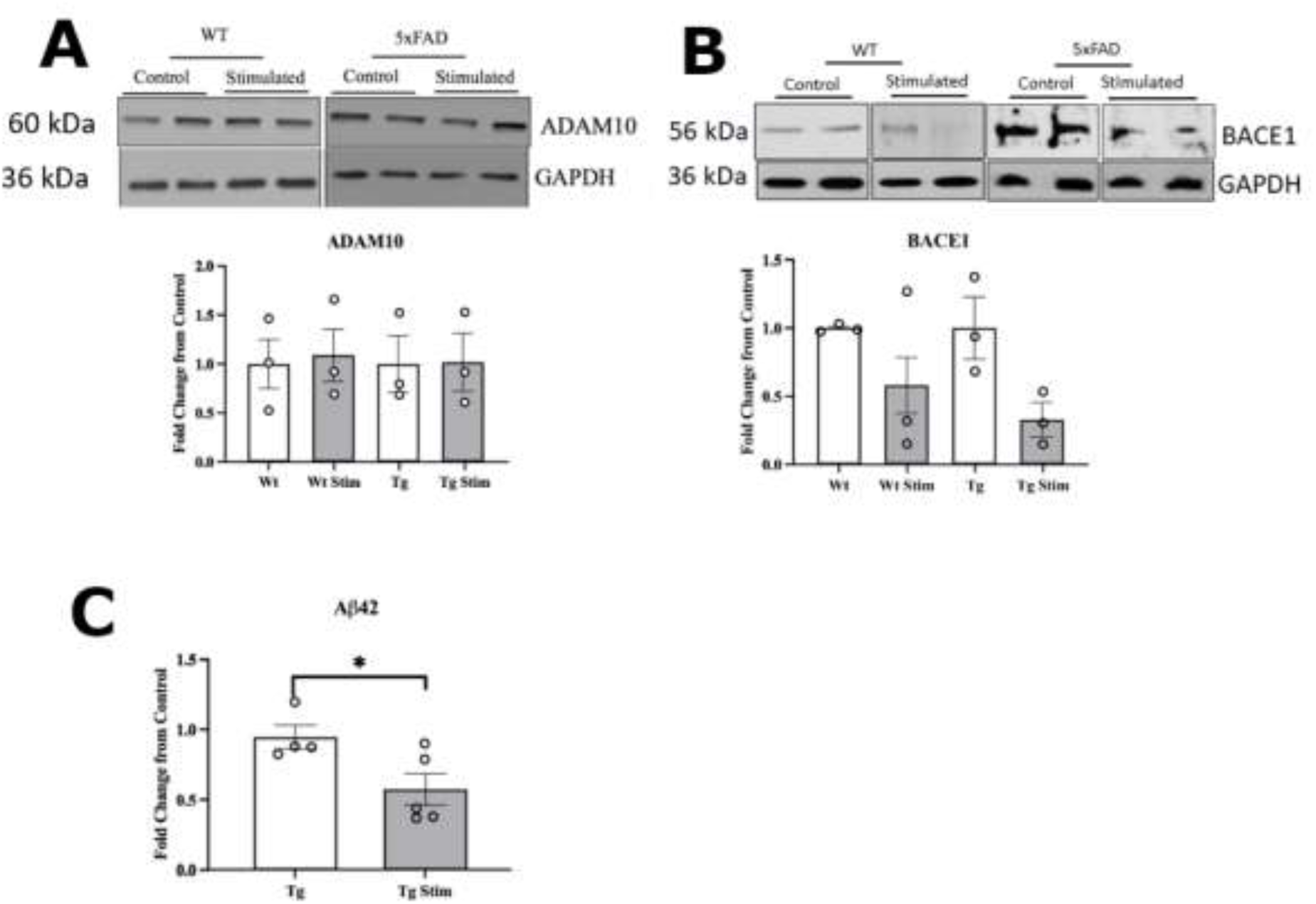
Concentrations of the amyloid precursor protein metabolic pathway markers after stimulation. Proteins were immunoprecipitated from frontal cortex of WT and 5xFAD mice and immunoblotted using antibodies described in the Materials and Methods section. The upper panel represents the immunoblot of the listed biomarker protein. The lower panel represents optical density values normalized against GAPDH and expressed as a percentage of the control group. Data is represented as mean per group ± SEM. *n* = 3 animals per group. (A) Immunoblot of ADAM10 at 60 kDa. Both groups showed no change in expression level. (B) Immunoblot of BACE1 at 56kDa. Both groups showed significant reduced expression with stimulation. WT t(4) = 2.04, p < 0.11. 5xFAD t(4) = 2.56, p < 0.06. *n* = 3 animals per group. (C) ELISA-measured concentration of Aβ_42_ showed significant decrease in stimulated 5xFAD mice frontal cortex compared to unstimulated controls. 5xFAD t(6) = 12.77, p < 0.001. *n* = 4-5. *** represents p < 0.001. Note: The TrkB blot (Fig 4A) was reprobed for ADAM10 (Fig 5A) whereas the TrkA blot (Fig 3A) was reporobed for BACE1 (Fig 5B).

### Hippocampal markers

The hippocampi of mice were also probed (Figure 6A). The projections from the basal forebrain densely target the cortical mantle, but only sparsely target the hippocampus, which received analogous innervation from the medial septum which was not stimulated. TrkA increases were found, and were significant in the 5xFAD groups, as shown in Figure 6B (5xFAD t(4) = 55.4, p < 0.0001). Reductions in BACE1 were also found in both genotypes and were significant in the wild-type genotype (WT t(4) = 12.77, p < 0.0003; 5xFAD t(4) = 1.23, p < 0.287; Figure 6C).

**Figure 6.**
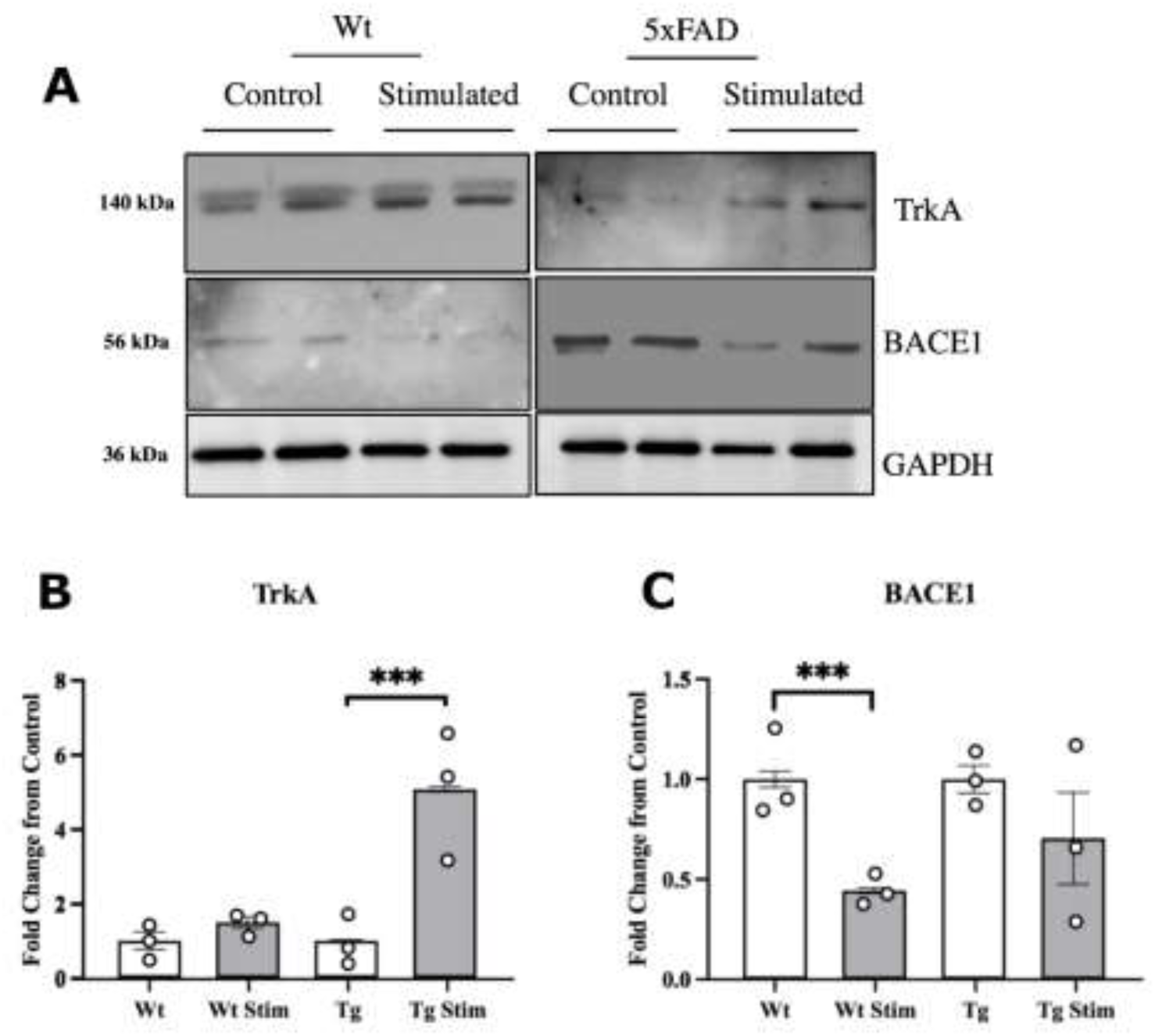
Neurotrophic and amyloid precursor protein metabolism markers in the hippocampus after nbM stimulation. Proteins were immunoblotted using antibodies described in the Materials and Methods section. A. The upper panel represents the immunoblot of the listed biomarker protein. The lower panel represents optical density values normalized against GAPDH and expressed as a percentage of the control group. Data is represented as mean per group ± SEM. *n* = 3 animals per group. C. Immunoblot of NGF’s receptor TrkA. Both mouse genotypes showed increases with stimulation and 5xFAD reached significance. 5xFAD t(4) = 55.4, p < 0.0001. D. Immunoblot of BACE1 at 56kDa. Both groups showed reduced expression with stimulation and WT reaching significance. WT t(4) = 12.77, p < 0.0003; 5xFAD t(4) = 1.23, p < 0.287. *n* = 3 animals per group.

## Discussion

The present study was conducted to assess the effects of non-concurrent intermittent stimulation of the basal forebrain, which contains cholinergic projection neurons, on brain pathways relevant in Alzheiemer’s dementia. To accomplish this, we stimulated mice that produce a high Aβ_42_ burden and display cognitive deficits. We used the Morris water maze to test stimulation impacts on visuospatial memory. Western blots and ELISAs allowed us to quantify the expression of neurotrophic and β-amyloidogenic markers of interest between treated and control groups.

The cognitive enhancement findings in this study are consistent with previous NGF and water maze related studies(Conner et al., 2009; De Rosa et al., 2005; Janis et al., 1997; Pham et al., 1999). We predicted NGF would be released during stimulation based on prior work(Hotta et al., 2009). Arguably, our experiment used endogenous acetylcholine pathways(Zaborszky et al., 2012) to trigger the creation of NGF in cortical receiving zones. Although the basal forebrain projects to the cortical mantle while the water maze more classically tests hippocampal function, it should be noted that immunotoxic lesioning of cholinergic neurons in the basal forebrain severely inhibits Morris water maze performance, while lesioning the hippocampal-projecting medial septum neurons has much less of a detrimental effect on performance(Berger-Sweeney et al., 1994; Mandel & Thal, 1988). Additionally, the basal forebrain is more of a complex continuum than completely segregated neuronal populations, so a hippocampal intensive task could still be relevant to a basal forebrain stimulation(Brashear et al., 1986; Carlsen et al., 1985; Zaborszky et al., 1991; Záborszky et al., 1986). Therefore, we believe the water maze to be an acceptable and even optimal behavioral experiment for testing basal forebrain-induced cognitive enhancement particularly because it uses a mildly aversive stimulus, cold water, to motivate escape behavior and thus constitutes a behavior with a strong reinforcement incentive.

Comparable studies to ours are fairly limited due to the advanced age and long-term treatment periods of our mouse groups as well as lab variability in pool size and average latency times (Gee et al., 2020; Schneider et al., 2014). Because our testing facility water maze is intended to also work with rats, its pool surface area is almost double the size of mouse-only studies making our task more difficult for mice. To date, there are two similar DBS studies worth comparing to this study. 3xTg AD mice were stimulated in the entorhinal cortex with tungsten wire electrodes which has been shown to increase neurogenesis in the dentate gyrus(Mann et al., 2018). Their stimulation pattern lasted for 7-hrs per day over the course of 25-days at a current half the amperage of this study and twice the frequency used. These mice demonstrated improvement in the probe test, but not in the 3-day acquisition period of their water maze(Mann et al., 2018). Aged APP/PS1 AD mice have been implanted with stainless steel electrodes in the basal forebrain and stimulating for various lengths of time and frequencies at a 10-fold stronger current than our study(Huang et al., 2019). They expectedly demonstrated that escape latencies in the water maze were best improved in the earliest implanted group at 4-mo old. Because the radius of the stimulation activating function increases in relation to the square root of the current(McIntyre et al., 2004; Stoney et al., 1968), the much higher 1mA current used would triple the stimulated tissue radius at the site of implantation. In other work, neither unstimulated 5xFAD nor wild type littermates demonstrate learning curves after the 5th day of the water maze(Schneider et al., 2014), similar to both of our non-treated mouse groups.

Our primary planned neurotrophic biomarker was TrkA. The mNGF receptor, TrkA, is a membrane-bound receptor that predicts cognitive loss, interacts with the amyloid precursor protein, and reports on the cholinergic system integrity(Boissiere et al., 1997; Matrone et al., 2011). We show here that our stimulation increases TrkA in both wild-type and 5xFAD mice in the cortex and also the hippocampus of 5xFAD mice. We believe this hippocampal effect may have occurred for one of a few reasons. The classic cholinergic subregions in the basal forebrain (Ch1-Ch4) are not entirely segregated. For example, some medial septum efferent fibers travel to the basal forebrain(Fujishiro et al., 2006; Richardson & DeLong, 1988). The stimulated region could also include any fiber, anterograde or retrograde, traveling through or nearby the basal forebrain and on their way to the hippocampus or medial septum.

The magnitude of the behavioral impacts of stimulation, and strong TrkA protein expression upregulation, motivated our look at brain derived neurotrophic pathway measures because it is also low in Alzheimer’s dementia post-mortem brains(Allen et al., 1999; Ferrer et al., 1999). Our finding, that TrkB is also upregulated, suggests the stimulation may use an intermediary messenger, a serine protease, that serves to cleave precursor forms of NGF and BDNF into mature forms(Bruno & Cuello, 2006; Gray & Ellis, 2008). This testable hypothesis has the benefit of one mechanism being caused by basal forebrain stimulation leading to activation of both neurotrophin receptors.

The evidence for direct activation of the neurotrophin receptors TrkA and TrkB was weaker in this study than the evidence receptor levels were high. The decision to wait over 24 hours after the last stimulation session before euthanasia and tissue analysis was made specifically to avoid acute impacts of stimulation, which fade within a few hours of activation(Kaplan et al., 1991; Soppet et al., 1991). One day later, normal receptor phosphorylation and neurotrophin levels would be expected. It may be the case that stimulation induced higher levels of mNGF and mBDNF, and led to phosphorylation of both their receptors, but those effects were largely absent by the 24 hour time point. A study directly comparing these same markers within an hour of finishing a stimulation session, and 24 hours later, could address this concern. Application of nicotine to PC12 cultures does result in upregulation of TrkA expression dependent on activation of α7 nicotinic receptors (Jonnala et al., 2002). This nicotine-induced expression has also been demonstrated in vivo(Formaggio et al., 2010; C. M. Hernandez & Terry, 2005). For these reasons we think the changes in TrkA expression are caused by activation of the cholinergic pathways, which activate nicotinic receptors, which induce an NGF response that leads to upregulation of TrkA. Similar relations are noted with nicotine and BDNF(Machaalani & Chen, 2018). Given the prevailing role of the nicotinic α7 receptor in both BDNF and NGF pathways, we hypothesize a common mediator of these effects: changes in extracellular serine protease activity triggered by α7 nicotinic receptor activation led to cleavage of precursor NGF and BDNF into mature form. We do not have evidence to rule out the nicotinic α7 receptor causing direct release of neurotrophins, however.

Our primary β-amyloidogenic marker was BACE1 since it is both the rate limiting step and necessary for the cleavage and subsequent production of Aβ_42_(Lahiri et al., 2014). Various cellular mechanisms have been found to regulate expression and activity of BACE1. In regards to the cholinergic system, p75^NTR^ activation leads to nSMase activation of ceramide which stabilizes BACE1’s active β-secretase site and increases its expression(Puglielli et al., 2003). Mice with a p75^NTR^ -/- genotype demonstrate a half-fold reduction in both BACE1 expression as well as overall Aβ_42_ burden which is striking given that the p75^NTR^ containing neurons are only the cholinergic neurons, which are significant a minority of the neuropil which otherwise has fairly ubiquitous BACE1 expression(Costantini et al., 2005). Ceramide elevation also inhibits mNGF-induced TrkA signaling, while TrkA, conversely, inhibits nSMase and ceramide production, creating a delicate balance between the proliferation and apoptosis pathways(MacPhee & Barker, 1999; Plo et al., 2004). We believe the decreased BACE1 signaling induced by basal forebrain stimulation is likely due to TrkA and TrkB-mediated PI3K/PKC signaling cascades that inhibit nSMase ceramide production, a testable hypothesis. Although BACE1 trended down with stimulation, it did not reach significance. Perhaps the reduction in overall Aβ_42_ burden merely occurred because it registered a cumulative signature of BACE1 expression changes.

## Acknowledgments

Founders for the 5xFAD mice were graciously provided by Raghavan Raju.

